# Neprilysin-sensitive amyloidogenic Aβ versus IDE-sensitive soluble Aβ: a probable mechanistic cause for sporadic Alzheimer’s disease

**DOI:** 10.1101/2021.08.22.457281

**Authors:** Hiroki Sasaguri, Risa Takamura, Naoto Watamura, Naomasa Kakiya, Toshio Ohshima, Ryo Fujioka, Naomi Yamazaki, Misaki Sekiguchi, Kaori Iwata, Yukio Matsuba, Shoko Hashimoto, Satoshi Tsubuki, Takashi Saito, Nobuhisa Iwata, Takaomi C. Saido

**Affiliations:** Laboratory for Proteolytic Neuroscience, RIKEN Center for Brain Science, 2-1 Hirosawa, Wako, Saitama 351-0198, Japan; Laboratory for Molecular Brain Science, Department of Life Science and Medical Bioscience, Waseda University, Shinjuku, Tokyo 162-8480, Japan; Department of Genome-based Drug Discovery & Unit for Brain Research, Graduate School of Biomedical Sciences, Nagasaki University, Nagasaki, 852-8521, Japan; Department of Neurocognitive Science, Institute of Brain Science, Nagoya City University Graduate School of Medical Sciences, Nagoya, Aichi 467-8601, Japan; Department of Neuroscience and Pathobiology, Research Institute of Environmental Medicine, Nagoya University, Nagoya, Aichi 464-8601, Japan

## Abstract

Neprilysin (NEP) and insulin-degrading enzyme (IDE) are considered the two major catabolic enzymes that degrade amyloid β peptide (Aβ), the primary cause of Alzheimer’s disease (AD). However, their roles in Aβ metabolism *in vivo* have never been compared in an impartial and side-by-side manner. Here, we crossbred single *App* knock-in mice with NEP (*Mme*) KO mice and with IDE (*Ide*) KO mice to generate double mutants that were analyzed for their biochemical and Aβ pathology properties. We found that NEP is responsible for the metabolism of amyloidogenic insoluble Aβ whereas IDE affects soluble Aβ. A deficiency of NEP, but not of IDE, augmented the formation of Aβ plaques, dystrophic neurites, and astrocytic and microglial activation, all of which are key pathological events in the development of AD. In addition, a deficiency of NEP had no significant impact on the levels of various neuropeptides (somatostatin, substance P, cholecystokinin, and neuropeptide Y), well known to be *in vitro* substrates for NEP, presumably because NEP is expressed in secretory vesicles and on the presynaptic membranes of excitatory neurons while most if not all neuropeptides are secreted from inhibitory neurons. This argues against the concern that NEP up-regulation for treatment of preclinical AD would reduce the levels of these neuropeptides. These findings indicate that NEP relatively selectively degrades Aβ in the brain. Whereas familial AD (FAD) is unambiguously caused by an increased anabolism of Aβ, and of Aβ _42_ and Aβ _43_ in particular, the anabolism of Aβ appears unaffected before its deposition in the brain that subsequently leads to the onset of sporadic AD (SAD). These observations thus suggest that NEP-sensitive amyloidogenic Aβ likely plays a primary pathogenic role in the etiology of SAD. Our findings are consistent with the aging-dependent decline of NEP expression in human brain and with recent genome-wide association studies (GWAS) indicating that variants of the gene encoding NEP (*MME*) are associated with the risk of SAD development. Taken together, our results imply that the aging-associated decrease in NEP expression is a primary cause of SAD and could thus be a target for the treatment of preclinical AD once other factors such as apolipoprotein E genotypes have also been considered.

## Introduction

Alzheimer’s disease (AD) is the major cause of dementia that deprives patients of their quality of life and dignity as the disease progresses. A large body of pathological and genetic evidence has established that the deposition of amyloid β peptide (Aβ) in the brain serves as a primary cause of this disorder and precedes the onset of fully developed AD by more than two decades ^1,2^. While familial AD (FAD) is unambiguously caused by the increased anabolism of Aβ (in particular Aβ _42_ and Aβ _43_), the mechanisms underlying Aβ accumulation in the etiology of sporadic AD (SAD) remain elusive. Because proteostasis is principally governed by the balance between anabolism and catabolism, and given that the anabolism of Aβ appears unaffected prior to the Aβ deposition that leads to SAD development, a plausible candidate cause of SAD is a decrease in Aβ catabolism ^3 4^. This notion led us to identify neprilysin (NEP) as a major Aβ -degrading enzyme *in vivo*, ^5,6^ while the group led by Selkoe identified insulin-degrading enzyme (IDE) as a contributing factor ^7-9^. From these findings it became commonplace for expression levels of these proteases to be quantified when examining the anabolism, clearance, or changes in Aβ *in vitro* or *in vivo* to ensure that catabolic mechanisms remained unaltered. However, as the quantitative and pathophysiological roles of NEP and IDE in *in vivo* Aβ catabolism have never been studied in an unbiased and side-by-side manner, we address this point in a series of animal experiments presented here.

To begin with, we make a clear distinction between two types of Aβ that are present in the brains of mutant mouse models of AD. The first can be solubilized by Tris-saline (TS) after homogenization and ultracentrifugation and is referred to as soluble Aβ ; the second requires a denaturing reagent, guanidine hydrochloride (GuHCl), for solubilization and is thus termed insoluble Aβ. This second type enhances Aβ amyloidosis *in vivo* and is referred to as being ‘amyloidogenic’ (see **Results**). In the present study, we used *Mme* KO mice^10^, which are deficient in NEP expression, and *Ide* KO mice^8^, which are deficient in IDE expression, and crossbred them with *App* knock-in mice, *App*^*NL-F/NL-F*^ that overproduce wild-type human Aβ _1-42_ without overexpressing amyloid precursor protein (APP) or APP/presenilin 1 (PS1) ^11,12^. The effect of NEP deficiency and that of IDE deficiency on AD-related pathological features were surprisingly different from each other in these animals. NEP deficiency resulted in a selective, significant increase in insoluble (amyloidogenic) Aβ but not in soluble Aβ as confirmed in biochemical and pathology analyses. On the other hand, a deficiency in IDE led to a specific increase in soluble Aβ _40_. Consequently, a NEP deficiency exacerbated Aβ amyloidosis and was associated with dystrophic neurites, loss of synaptic markers and inflammatory responses, whereas IDE deficiency exerted essentially no pathological effects. These finding support the notion that NEP is a primary Aβ -degrading enzyme both under physiological and pathological conditions.

When we showed that gene therapy utilizing NEP activity attenuated Aβ pathology in APP-transgenic mice^13-15^ and that the neuropeptide somatostatin regulated neuronal Aβ levels *in vitro* and *in vivo*^*16-19*^, interest within the academic and pharmaceutical industry fields rose significantly given the prospects of applying pharmacological intervention via the somatostatin-NEP pathway to suppress Aβ pathology ^20-23^ and thus inhibit or delay the progression of preclinical AD. This concept was consistent with the aging-dependent reduction and oxidative inactivation of neprilysin,^24-27^ along with the disappearance of somatostatin with aging and in AD^28,29^, all of which would lead to accelerated Aβ deposition. Finally, while recent GWAS indicated an association of *MME* gene variants with the incidence of AD ^30 31^, to our knowledge there has been no such relationship reported between the *IDE* gene and risk of AD.

Some concerns have been raised in the scientific literature about modulating NEP activity to treat preclinical AD due to postulated effects on the levels of various other neuropeptides present in the central nervous system (CNS): NEP was originally described as a peptidase that degraded neuropeptides *in vitro* by biochemical means^32-34^. We thus examined the levels of representative CNS neuropeptides such as somatostatin, substance P, cholecystokinin, and neuropeptide Y in *Mme* KO mouse brains and unexpectedly found that a deficiency of NEP did not elevate these neuropeptides in the CNS, but did result in a doubling of endogenous Aβ levels^6^. This observation agreed with an early report showing that enkephalin levels remained unchanged in the cerebral cortices of *Mme* KO mice ^35^, and occurs presumably because NEP degrades Aβ in secretory vesicles and at presynaptic membranes of excitatory neurons^26,36,37^. Furthermore, the aforementioned neuropeptides are generally produced and secreted from inhibitory neurons^38-41^, thereby accounting for discrepancies between *in vivo* and *in vitro* observations. Now that we are aware of the involvement of somatostatin receptor subtypes 1 and 4 in neprilysin-catalyzed Aβ regulation ^17^, our findings together point to a G protein-coupled receptor (GPCR)-based strategy for the generation of low molecular weight, disease-modifying medications to treat preclinical AD, which would be much less expensive and likely to be safer than immunotherapies.

## Results

### Aβ profiles in the brains of *Mme* and *Ide* KO mice

To evaluate the roles of NEP and IDE in the physiological degradation of Aβ *in vivo*, we compared Aβ profiles in the brains of single *Mme* KO (NEP-deficient) and *Ide* KO (IDE-deficient) mice by Enzyme-Linked Immunosorbent Assay (ELISA) (**Figure 1**). The *Mme* KO mice showed 1.5-, 1.8- and 1.6-fold increases in TS-soluble Aβ_42_, GuHCl-soluble Aβ_40_ and Aβ_42_, respectively (**Figure 1a,b,d,e**). In contrast, the quantity of TS-soluble Aβ_40_ and Aβ_42_ significantly increased by 5.6- and 2.1-fold, respectively, in *Ide* KO mice (**Figure 1a,d**) whereas the GuHCl-soluble Aβ_40_ and Aβ_42_ fractions remained unchanged (**Figure 1b,e**). It is noteworthy that the quantity of GuHCl-soluble Aβ was approximately 10 times greater than that of TS-soluble Aβ in wild-type brains. The total amount of Aβ_40_ and Aβ_42_ increased significantly in *Mme* KO mice, whereas the total Aβ_40_ content in *Ide*-KO mice increased only marginally (**Figure 1c, f**). For the Aβ_42_/Aβ_40_ ratio, both mouse lines showed similar profiles (**Figure 1g-i**). A deficiency of NEP or IDE decreased the Aβ_42_/Aβ_40_ ratio in TS-soluble fractions but did not affect those in GuHCl and total fractions. These results indicate that IDE mainly targets TS-soluble Aβ, whereas NEP can degrade insoluble Aβ species that account for the majority of CNS Aβ accumulation *in vivo*.

**Fig. 1.**
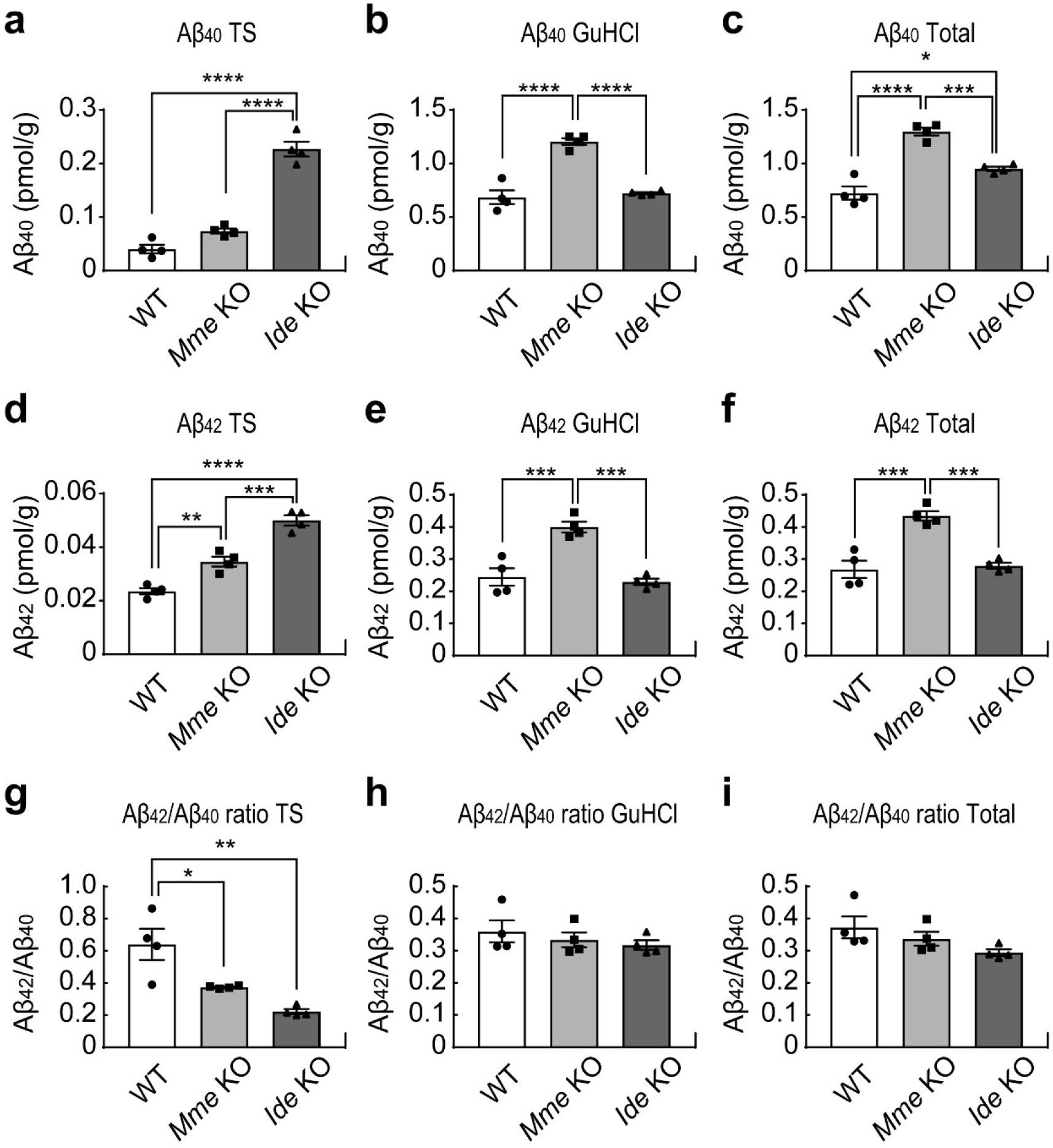
Aβ in *Mme* KO and *Ide* KO mice. **a-f**, Measurement of Aβ_42_ (**a-c**) or Aβ_40_ (**d-f**) levels by ELISA in Tris-buffered saline (TS) fractions (**a, d, g**), guanidine hydrochloride (GuHCl) fractions (**b, e, h**) and total fractions (**c, f, i**) from the brains of 12-month-old wild type (WT), *membrane metallo endopeptidase* (*Mme*, encoding NEP)-knock-out (KO) and *insulin degrading enzyme* (*Ide*) KO mice (*n* = 4). One-way ANOVA followed by Tukey’s multiple comparisons test. \**p* < 0.05, \*\**p* < 0.01, \*\*\**p*< 0.001, \*\*\*\**p* < 0.0001. **g-i**, Aβ_42_/ Aβ_40_ ratios in TS, GuHCl and total fractions prepared from brains of 12-month-old WT, *Mme* KO, and *Ide* KO mice (*n* = 4).

### Processing of APP in *Mme* and *Ide* KO mice

We next examined APP processing in *Mme* and *Ide* KO mice. APP is first cleaved either by α-secretase or β-secretase (a type I transmembrane aspartic protease also termed β-site APP-cleaving enzyme 1 or BACE1) to produce the C terminal fragments (CTFs), CTF-α and CTF-β, respectively ^42^. There was no significant difference in the amounts of full-length APP or APP CTFs in wild-type mice, *Mme* or *Ide* KO mice (**Figure 2a**), indicating that a deficiency of NEP or IDE did not affect the production or α/β-secretase-mediated processing of APP. The amount of NEP protein in *Ide* KO mice was, however, significantly increased (**Figure 2a,b**). Because the APP intracellular domain (AICD) transcriptionally regulates *Mme* gene expression ^43-47^ and given that both IDE and AICD are mainly localized in the cytoplasm, it is possible that the increased NEP level in *Ide* KO mice was caused by an increase in AICD. In contrast, the level of IDE was markedly decreased in *Mme* KO mice (**Figure 2a,c**); while the mechanism for this remains unclear, it may account for the elevation of TS-soluble Aβ in *Mme* KO mice in a manner similar to that of *Ide* KO mice (**Figure 1a,d,g**).

**Fig. 2.**
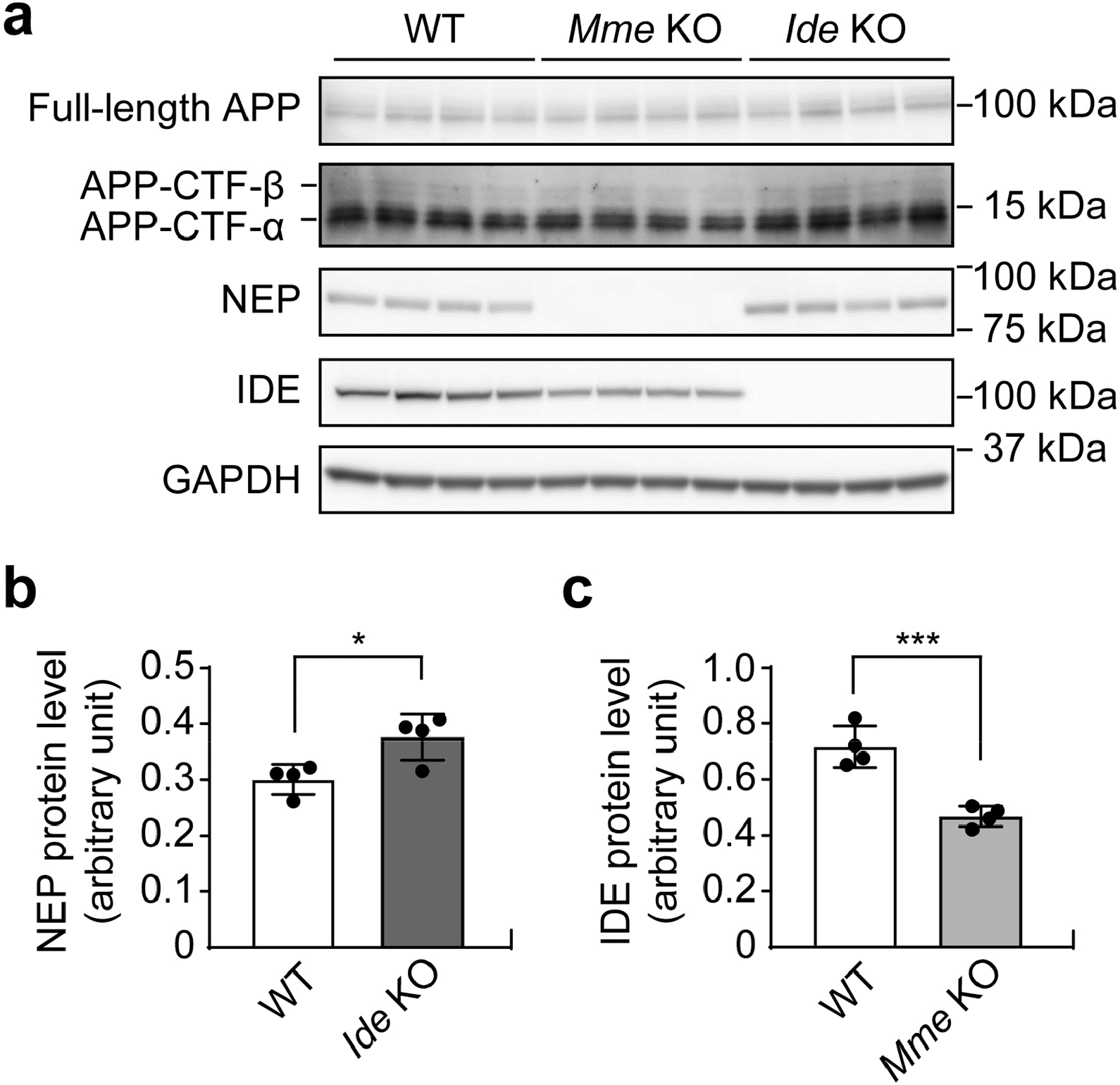
Processing of APP in *Mme* KO and *Ide* KO mice. **a**, Western blot of full-length (FL) APP, APP C terminal fragments (CTF), NEP, IDE, and GAPDH in the cortices of 12-month-old WT, *Mme* KO and *Ide* KO mice (*n* = 4). **b, c**, Quantifications of NEP (**b**) and IDE (**c**) in the cortices of 12-month-old WT and *Mme* KO or *Ide* KO mice (*n* = 4). Student’s *t*-test. *p>0.05, ***p>0.001. A deficiency of NEP or IDE exerted no effect on APP expression or β-cleavage activity, confirming that changes in Aβ levels were due to alterations in Aβ degradation. The expression of NEP was increased in *Ide* KO mice, while the expression of IDE was decreased in *Mme* KO mice.

### Effects of NEP and IDE deficiency on the Aβ amyloidosis in *App* knock-in mice

To elucidate the roles of NEP and IDE in the pathophysiology of AD, we crossbred *Mme* and *Ide* KO mice with *App*^*NL-F/NL-F*^ mice^11^, the latter of which harbor the Swedish (KM670/671NL)^48^ and Beyreuther/Iberian (I716F) ^49^ mutations in the endogenous mouse *App* gene, and recapitulate typical Aβ pathology and neuroinflammation in brain tissue from approximately 8 months of age. We used ELISA to quantify TS- and GuHCl-soluble Aβ in 12-month-old mice and found that a deficiency of NEP or IDE in *App*^*NL-F/NL-F*^ mice changed the Aβ profiles to patterns similar to those seen in the single *Mme* and *Ide* KO mice (**Figures 1 and 3**). These findings indicate that NEP and IDE play analogous roles both under physiological and pathological conditions. *App*^*NL-F/NL-F*^ X *Mme* KO mice showed a significant increase of GuHCl-soluble Aβ_40_ and Aβ_42_ (**Figure 3a,b,d,e**), whereas *App*^*NL-F/NL-F*^ X *Ide* KO mice exhibited an increase in TS-soluble Aβ_40_ and Aβ_42_, although the increase in TS-soluble Aβ_42_ was not statistically significant. The total amount of Aβ_40_ and Aβ_42_ increased significantly only in the *App*^*NL-F/NL-F*^ X *Mme* KO mice (**Figure 3c, f**). This observation is consistent with a previous report showing that IDE degrades Aβ monomer but not oligomer species^50^. A deficiency of IDE in *App*^*NL-F/NL-F*^ mice tended to decrease the Aβ_42_/Aβ_40_ ratio in TS-soluble fractions although this was not statistically significant (**Figure 3g-i**). In contrast, the Aβ_42_/Aβ_40_ ratio in GuHCl fractions and total fractions was not affected in either of these lines. These results confirm that IDE primarily targets TS-soluble Aβ, whereas NEP degrades more insoluble Aβ species as well as under pathological conditions. Gel filtration analyses of each soluble fraction indicated that deficiencies both in NEP and IDE induced the generation primarily of the Aβ _42_ monomer along with relatively small quantities of dimer, trimer and tetramer species (**Supplementary Figure 1**), the profiles of which did not differ between *Mme* KO and *Ide* KO mice crossbred with *App* knock-in mice. These observations indicate that soluble Aβ oligomer species constitute a very minor proportion of overall Aβ *in vivo* even under pathological conditions.

**Fig. 3.**
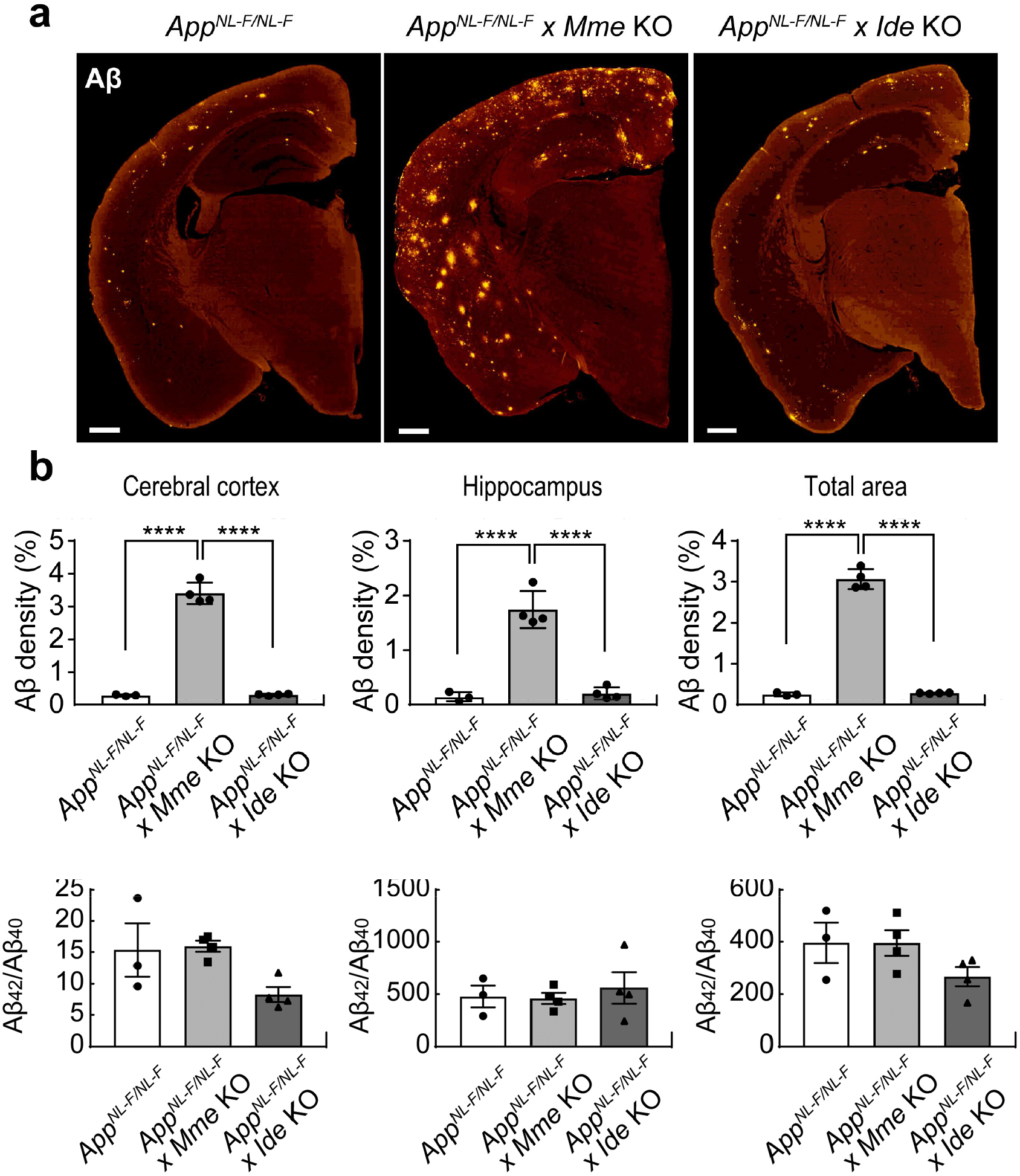
Aβ in *App*^*NL-F/NL-F*^ x *Mme* KO and *App*^*NL-F/NL-F*^ x *Ide* KO mice. **a-f**, Measurement of Aβ_42_ (**a-c**) or Aβ_40_ (**d-f**) levels by ELISA in TS fractions (**a, d, g**), GuHCl fractions (**b, e, h**), and total fractions (**c, f, i**) prepared from brains of 12-month-old *App*^*NL-F/NL-F*^, *App*^*NL-F/NL-F*^ x *Mme* KO and *App*^*NL-F/NL-F*^ x *Ide* KO mice (*n* = 3∼4). **g-i**, Aβ42/ Aβ40 ratio in brains of 12-month-old *App*^*NL-F/NL-F*^, *App*^*NL-F/NL-F*^ x *Mme* KO and *App*^*NL-F/NL-F*^ x *Ide* KO mice (*n* = 3∼4). One-way ANOVA followed by Tukey’s multiple comparisons test. *p < 0.05, **p < 0.01, ***p < 0.001. NEP mainly degrades GuHCl-soluble (TS-insoluble) Aβ species, whereas IDE mainly targets TS-soluble Aβ.

### Neuropathology of *App*^*NL-F/NL-F*^ X *Mme* KO and *App*^*NL-F/NL-F*^ X *Ide* KO mice

Next, we evaluated the roles of NEP and IDE in the AD pathology expressed by our animal models. We performed immunohistochemical analyses of Aβ in brain slices and found that a lack of IDE did not affect Aβ plaque load, whereas a deficiency of NEP significantly exacerbated the Aβ pathology both in the cortex and hippocampus (**Figure 4a,b**). Neuroinflammation induced by activated astrocytes and microglia surrounding amyloid plaques manifests as one of the main pathological features in AD patients and AD mouse models^11,30^. We analyzed the neuroinflammatory status of *App*^*NL-F/NL-F*^ X *Mme* KO and *App*^*NL-F/NL-F*^ X *Ide* KO mice by immunofluorescence using antibodies against astrocytes (GFAP) and microglia (Iba1) (**Figure 5a-c**). *App*^*NL-F/NL-F*^ X *Mme* KO mice showed increased astrocytosis and microgliosis in the cortex, whereas in *App*^*NL-F/NL-F*^ X *Ide* KO mice the extent of neuroinflammation was comparable to that of *App*^*NL-F/NL-F*^ mice. This finding indicates that neuroinflammation in AD is mainly induced by GuHCl-soluble Aβ rather than TS-soluble Aβ species. In addition, we detected similar losses of synaptophysin and PSD95 immunoreactivities in the vicinity of Aβ plaques compared to those observed in our previous AD mouse models^11^ (**Figure 5d,e**). Taken together, our results reveal that NEP mainly degrades Aβ species with low solubility and affecting AD-related neuropathology, while IDE targets soluble Aβ *in vivo* that does not affect such pathological features.

**Fig. 4.** Aβ deposition in *App*^*NL-F/NL-F*^ x *Mme* KO and *App*^*NL-F/NL-F*^ x *Ide* KO mice. **a**, Aβ deposition in *App*^*NL-F/NL-F*^ (left), *App*^*NL-F/NL-F*^ x *Mme* KO (middle), and *App*^*NL-F/NL-F*^ x *Ide* KO (right) mouse brains. Brain sections from 12-month-old mice were immunostained using antibody to N-terminal Aβ (N1D). Scale bars represent 500 µm. **b**, Area quantification of N1D immunostaining in the cortex (left), hippocampus (middle) and hemicephalia (right) of 12-month-old *App*^*NL-F/NL-F*^, *App*^*NL-F/NL-F*^ x *Mme* KO and *App*^*NL-F/NL-F*^ x *Ide* KO mice (*n* = 3∼4). One-way ANOVA followed by Tukey’s multiple comparisons test. ****p<0.0001.

**Fig. 5.**
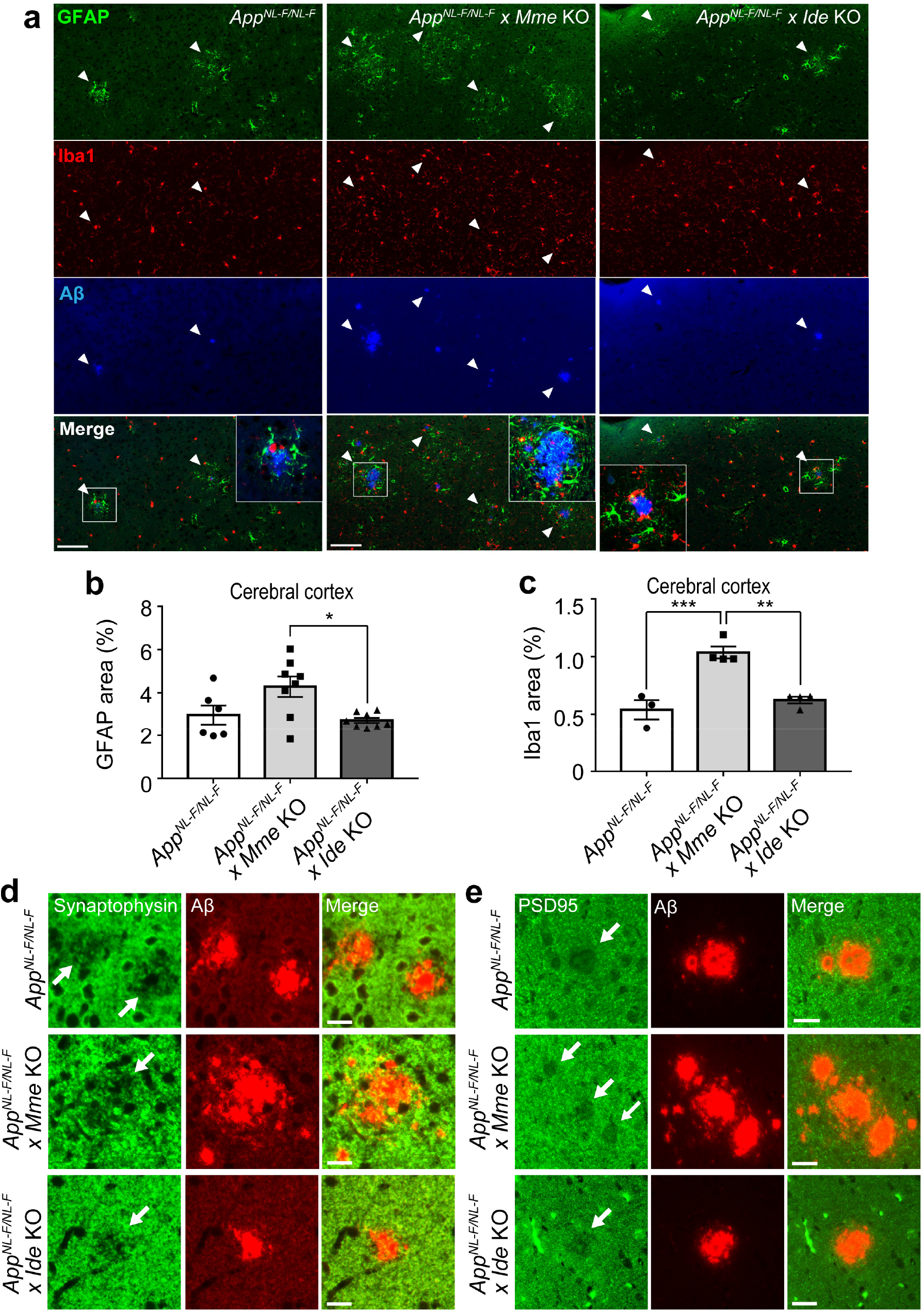
Neuroinflammation and synaptic alterations in *App*^*NL-F/NL-F*^ x *Mme* KO and *App*^*NL-F/NL-F*^x *Ide* KO mice. **a**, Astrocytosis (green) and microgliosis (red) in *App*^*NL-F/NL-F*^ (left), *App*^*NL-F/NL-F*^ x *Mme* KO (middle), and *App*^*NL-F/NL-F*^ x *Ide* KO (right) mouse brains. Brain sections from 12-month-old mice were immunostained using antibody to GFAP, Iba1, and N-terminal Aβ (N1D, blue). Scale bars represent 500 µm (low magnification) and 100 µm (high magnification). **b, c**, Area quantification of astrocytosis and microgliosis (**c**) in the cortices of 12-month-old *App*^*NL-F/NL-F*^, *App*^*NL-F/NL-F*^ x *Mme* KO and *App*^*NL-F/NL-F*^ x *Ide* KO mice (*n* = 3∼4). One-way ANOVA followed by Tukey’s multiple comparisons test. \**p*<0.05, \*\**p*<0.01, and \*\*\**p*<0.001. **d, e**, Synaptic alterations in the brains of 12-month-old *App*^*NL-F/NL-F*^, *App*^*NL-F/NL-F*^ x *Mme* KO and *App*^*NL-F/NL-F*^ x *Ide* KO mice. Double staining was performed using N1D (**d**) or 82E1 (**e**) antibodies with a presynaptic marker (antibody to synaptophysin, **d**) and with a postsynaptic marker (antibody to PSD95, **e**). Scale bars represent 10 μm.

### Neuropeptide profiles in *Mme* KO mice

Our results raised the possibility that NEP could be targeted – by increasing its expression or activity – to reduce GuHCl-soluble Aβ or amyloid plaques in preclinical AD. However, NEP has been reported to also degrade neuropeptides as well as Aβ in the CNS ^4,32^. To evaluate the effect of NEP on the neuropeptide levels *in vivo*, we performed immunohistochemistry against somatostatin, substance P, cholecystokinin, and neuropeptide Y in single *Mme*-KO mice (**Figure 6** and **Supplementary Figures 2-5**). We confirmed that a deficiency of NEP did not alter the levels of neuropeptides except for that of somatostatin in the subiculum, where a small but significant reduction was seen (**Figure 6**). In any case, NEP deficiency did not induce an increase in neuropeptide levels, suggesting that the pharmacological up-regulation of NEP activity to reduce brain Aβ would not induce side effects caused by reduced neuropeptide levels.

**Fig. 6.**
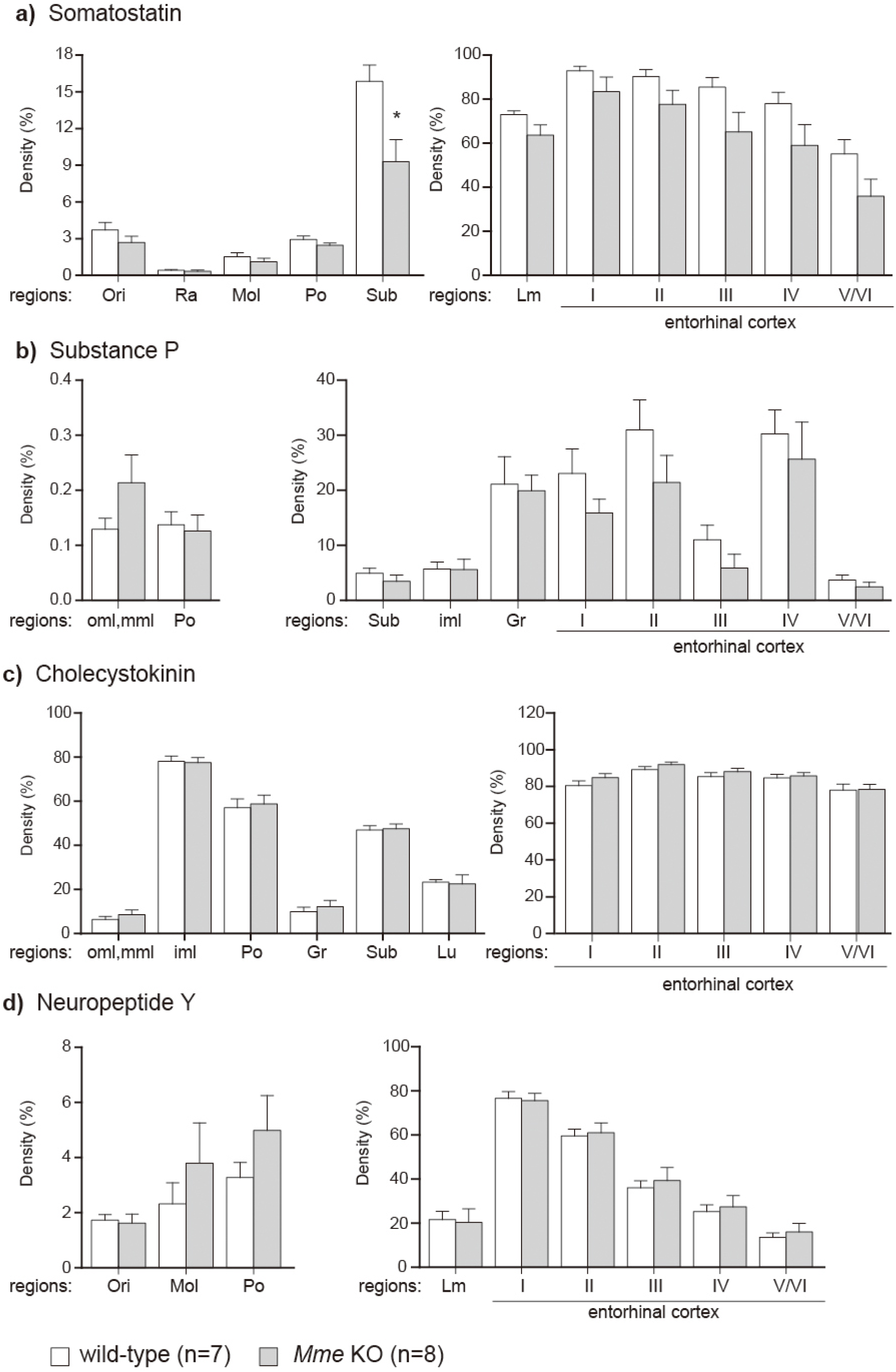
Quantification of immunohistochemical images for four neuropeptides (somatostatin, substance P, cholecystokinin, and neuropeptide Y) in wild-type and *Mme* KO mouse brains. Representative immunohistochemistry images are shown in Supplementary Figures 2-5. The open and gray columns indicate wild-type and *Mme* KO mice, respectively. Mice deficient in somatostatin precursor ^16^ were used as a negative control to confirm specificity of the staining. The abbreviations used are represented in Fig. 2S. *p < 0.05, significantly different from wild-type tissues.

### Memory impairment in single *Mme* KO mice

We considered that it would be interesting to compare levels of cognition between *App*^*NL-F/NL-F*^ X *Mme* KO and *App*^*NL-F/NL-F*^ X *Ide* KO mice as this might provide information on whether the NEP-sensitive or IDE-sensitive Aβ is more neurotoxic. We however found that single *Mme* KO mice exhibited significant memory impairment at 6 months of age as detected by a fear conditioning test (**Supplementary Figure 6**). This fact makes it difficult to assess the cognitive status of these double mutant mice in an unbiased manner, although the pathological features of the *Mme* KO mouse line (i.e., enhanced amyloidosis and neuroinflammation (**Figure 5**)), which were shown in a number of studies to be associated with memory impairment ^11,51-54^, suggest that NEP-sensitive amyloidogenic Aβ rather than IDE-sensitive soluble Aβ is principally responsible for the neurotoxicity present *in vivo*.

## Discussion

Two decades have passed since we demonstrated that NEP is one of the major physiological Aβ -degrading enzymes ^5,6^. A mathematical formulation, based on the rate constants for anabolism, catabolism, and clearance (to cerebral spinal fluid/plasma) of Aβ, indicated that NEP activity would account for around 50 % of all catabolism/clearance mechanisms given that a deficiency of NEP led to an approximately 2-fold increase in Aβ levels *in vivo* ^3^. It is notable that just a 50% increase in Aβ production results in early Aβ deposition in some cases of FAD and Down’s syndrome^2^, implying that a 50% reduction of NEP expression/activity, which increase endogenous brain Aβ levels 1.5-fold ^6^, can cause pathological Aβ deposition leading to AD development. Our present study showed that NEP is responsible not only for physiological Aβ metabolism but also for pathological Aβ metabolism. During these past decades, a large body of genetic, pathological and molecular biological evidence has established empirical roles for neuroinflammation in the pathogenesis of AD ^31,55-57^. Consistently, using single *App* knock-in mice we have shown that the NEP-sensitive amyloidogenic Aβ was closely associated with neuroinflammation (**Figure 5**). Given that recent GWAS outcomes implied a close association between *MME* gene variants and AD incidence ^30,31^, NEP now stands as one of the key players in AD pathogenesis that may serve as a druggable target for the treatment of preclinical AD.

We previously predicted the presence of a mechanism in the CNS whereby neuropeptide(s) control Aβ levels via the regulation of NEP activity ^4^. Indeed, we found that somatostatin activates NEP *in vitro* and *in vivo* and that a deficiency of somatostatin receptor subtypes 1 and 4 augmented Aβ deposition by diminishing NEP expression in the brain ^17^, indicating that specific agonist(s) or allosteric modulator(s) to such receptors may attenuate Aβ deposition in the preclinical stage. Because somatostatin receptor subtypes are G protein-coupled receptors (GPCRs)^17^, the selective up-regulation of NEP by pharmacological means targeting at such GPCR(s) may provide an effective, safe, and socioeconomically affordable disease-modifying treatment for preclinical AD patients. Such treatments may replace costly immunotherapy options ^58^ as GPCRs are the best targets for low molecular weight medications. This strategy is again consistent with the aging-dependent reduction and oxidative inactivation of neprilysin,^24-27^ along with the disappearance of somatostatin with aging and in AD^28,29^, all of which would lead to accelerated Aβ deposition.

Concerns however have been expressed in pharmaceutical industry about modifying NEP activity to reduce Aβ levels in the brain because NEP has been suggested to degrade neuropeptides *in vitro*^32-34^. Nonetheless, consistent with Saria et al., who showed that enkephalin levels remained unchanged in the cortices of *Mme* KO mice ^35^, a deficiency of NEP had no significant impact on somatostatin, substance P, cholecystokinin, or neuropeptide Y levels in our experiments (**Figure 6**). This is presumably because NEP degrades its substrate(s) inside secretory vesicles and on the presynaptic membrane of excitatory neurons^26,36,37^ and because inhibitory neurons in general secrete neuropeptides^38-41^, making it difficult for these neuropeptides to encounter NEP *in vivo*. In contrast, NEP appears to serve as a functional peptidase that degrades neuropeptide(s) in organs such as the heart and kidney^59,60^.

In conclusion, we emphasize that aging-associated down-regulation of NEP is likely a primary cause for SAD and that up-regulation of NEP expression or its activity in the CNS via somatostatin receptor activation should be considered as a strategy to reduce Aβ deposition in preclinical AD to either halt or delay onset of the disease. Our 3^rd^ generation mouse model of AD that accumulates wild-type human Aβ very rapidly^61^ will become a powerful tool for the primary *in vivo* screening of such medication candidates.

## Materials and Methods

### Animals

*Mme*-KO mice ^10^ were kindly provided by Craig Gerard, Harvard Medical School, USA. *Ide*-KO mice ^8^ were purchased from MMRRC at the University of California, Davis. The generation of *App*^*NL-F/NL-F*^ mice was described previously^11^. *Mme* and *Ide* KO mice were crossbred with *App*^*NL-F/NL-F*^ mice to generate *App*^*NL-F/NL-F*^ X *Mme* KO and *App*^*NL-F/NL-F*^ X *Ide* KO mice, respectively. All double mutant mice used in this study were homozygous for the *App* mutations and NEP or IDE deficiency. C57BL/6J mice were used as wild type controls. Both male and female mice were used for biochemical and immunohistochemical studies. All mice were bred and maintained in accordance with regulations for animal experiments promulgated by the RIKEN Center for Brain Science.

### Genotyping

Genomic DNA was extracted from mouse tails in lysis buffer (10 mM pH 8.5 Tris-HCl, 5 mM pH 8.0 EDTA, 0.2% SDS, 200 mM NaCl, 20 µg/ml proteinase K) and subjected to PCR. Primers used for genotyping were: 5′-TCCAAATGTGTCAGTTTCATAGCC-3′, 5′-GGCTATGACTCATGATGTCATAACAGG-3′, and 5′-GCCTATTCTTACCAAATATTCTCCCAG-3′ for *Mme* KO mice; 5′-ATAAACCCTCTTGCAGTTGCATC-3′, 5′-ACATACTTCCCAGAGCATAGGACG-3′, and 5′-CTAATGAAACTGGGAGGGTTGG-3′ for *Ide* KO mice; and 5′-ATCTCGGAAGTGAAGATG-3′, 5′-ATCTCGGAAGTGAATCTA-3′, 5′-TGTAGATGAGAACTTAAC-3′ and 5′-CGTATAATGTATGCTATACGAAG-3′ for *App*^*NL-F/NL-F*^ mice.

### Brain sample preparation

Mice were anesthetized with isoflurane and transcardially perfused with ice-cold PBS ^11^. Brains were extracted, maintained on ice, and dissected into two halves at the midline. For biochemical analyses, one hemisphere was divided into several parts including the hippocampus, and stored at -80 °C. For immunohistochemical analyses, the brain was fixed with 4% paraformaldehyde in PBS, incubated at 4 °C for 24 h and rinsed with PBS until paraffin processing.

### Western blotting

Mice brain tissues were homogenized in lysis buffer (50 mM Tris pH 7.6, 0.15 M NaCl and Complete protease inhibitor cocktail (Roche)) ^11^. Homogenates were incubated at 4 °C for 1 h, centrifuged at 15,000 rpm for 30 min, and the supernatants were collected. Equal amounts of proteins per lane were subjected to SDS-PAGE and transferred to PVDF or nitrocellulose membranes (Invitrogen). To detect APP-CTFs, delipidated samples were loaded and the membrane was boiled for 5 min in PBS before the next step. After washing and blocking at room temperature, the membranes were incubated at 4 °C overnight with primary antibodies against full-length APP (1:1,000, Millipore), APP-CTFs (1:1,000, Sigma-Aldrich), NEP (1:500, R&D Systems), IDE (1:1,000, Abcam) or against GAPDH as a loading control (HRP conjugated, 1:150,000, ProteinTech). The target protein on the membrane was visualized with ECL Select (GE Healthcare) and a Luminescent Image Analyzer LAS-3000 Mini (Fujifilm).

### Immunostaining

Paraffin-embedded mouse brain sections were subjected to deparaffinization and then antigen retrieval was performed by autoclave processing at 121 °C for 5 min ^11^. After inactivation of endogenous peroxidases using 0.3% H_2_O_2_ solution for 30 min, the sections were washed with TNT buffer (0.1 M Tris pH 7.5, 0.15 M NaCl, 0.05% Tween20), blocked for 30 min in TNB buffer (0.1 M Tris pH 7.5, 0.15 M NaCl) and incubated overnight at 4 °C in the same buffer with primary antibodies. The primary antibody dilution ratios were as follows: Aβ1-5 (N1D) ^62^ (1:200), GFAP (1:200, Millipore), Iba1 (1:200, Wako), synaptophysin (1:200, PROGEN), and PSD95 (1:50). Amyloid pathology was detected using biotinylated secondary antibody and tyramide signal amplification as described previously (Enya, 1999). Before mounting, the sections were treated when necessary with Hoechst33342 diluted in PBS. Data images were obtained using a NanoZoomer Digital Pathology C9600 (Hamamatsu Photonics) and EVOS M5000 Imaging System (Thermo Fisher scientific). Immunoreactive signals were quantified by Definiens Tissue Studio (Definiens).

### ELISA

Mouse cortical samples were homogenized in buffer A (50 mM Tris-HCl, pH 7.6, 150 mM NaCl and protease inhibitor cocktail) using a medical beads shocker ^11^. The homogenized samples were directed to centrifugation at 200,000×g for 20 min at 4 °C, and the supernatant was collected as a TS-soluble fraction. The pellet was loosened with buffer A, centrifuged at 200,000×g for 5 min at 4 °C, and then dissolved in 6 M Gu-HCl buffer. After incubation at room temperature for 1 h, the sample was sonicated at 25 °C for 1 min. Subsequently, the sample was centrifuged at 200,000×g for 20 min at 25°C and the supernatant collected as a GuHCl fraction. 100 µl of TS and GuHCl fractions were loaded onto 96-well plates and incubated at 4 °C overnight using the Aβ_40_ and Aβ_42_ ELISA kit (Wako) according to the manufacturer’s instructions.

### Statistics

All data are shown as the mean ± S.E.M. For comparisons between two groups, statistical analyses were conducted by Student’s *t*-test. For comparisons among three groups, one-way analysis of variance (ANOVA) was used followed by Tukey’s multiple comparisons test. These analyses were performed using GraphPad Prizm 8 software (GraphPad software). The levels of statistical significance were shown as P-values: * *P* < 0.05, ** *P* < 0.01, *** *P* < 0.001, and **** *P* < 0.0001.

## Author Contributions

HS, RT, TS, NI, and TCS designed the research plan. HS, RT, NW, NK, RF, NY, MS, KW, YM, and ST performed the experiments. HS, RT, NK, SH, ST, TS, NI, and TCS analyzed and interpreted data. HS, RT, NW, SH, TS, NI and TCS wrote the manuscript. HS, TO, TS, NI and TCS supervised the entire research progress.

## Acknowledgements

We thank Taisuke Tomita, University of Tokyo, for valuable discussion. We also thank Yukiko Nagai-Watanabe for secretarial work. This work was supported by AMED under Grant Number JP20dm0207001 (Brain Mapping by Integrated Neurotechnologies for Disease Studies (Brain/MINDS)) (TCS) and JSPS KAKENHI Grant Number JP18K07402 (HS).

## Conflicts of interest

The authors have no conflicts of interest to declare.

## Supplementary Information

**Fig. 1S.**
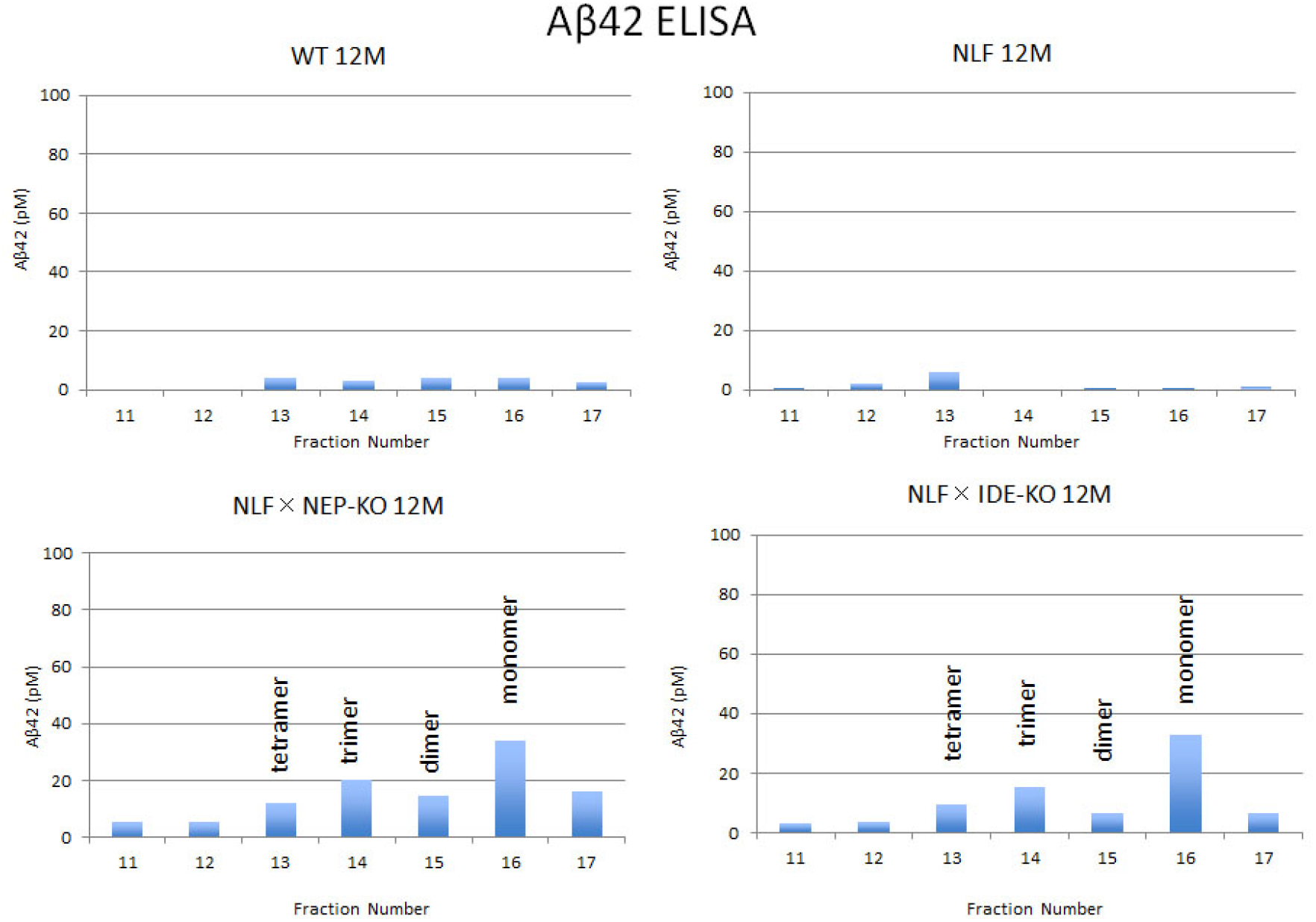
Molecular sieving analysis of Aβ_42_ by gel filtration of TS-soluble extracts. TS-soluble fractions from brain extracts were subjected to gel filtration analysis for detection of Aβ monomer and oligomers. *App*^*NL-F*^ mice deficient in NEP and IDE exhibited similar quantities of monomer and oligomers (i.e., from the dimer to tetramer). Aβ_40_ levels were below the detection limit in all fractions (data not shown).

**Fig. 2S.**
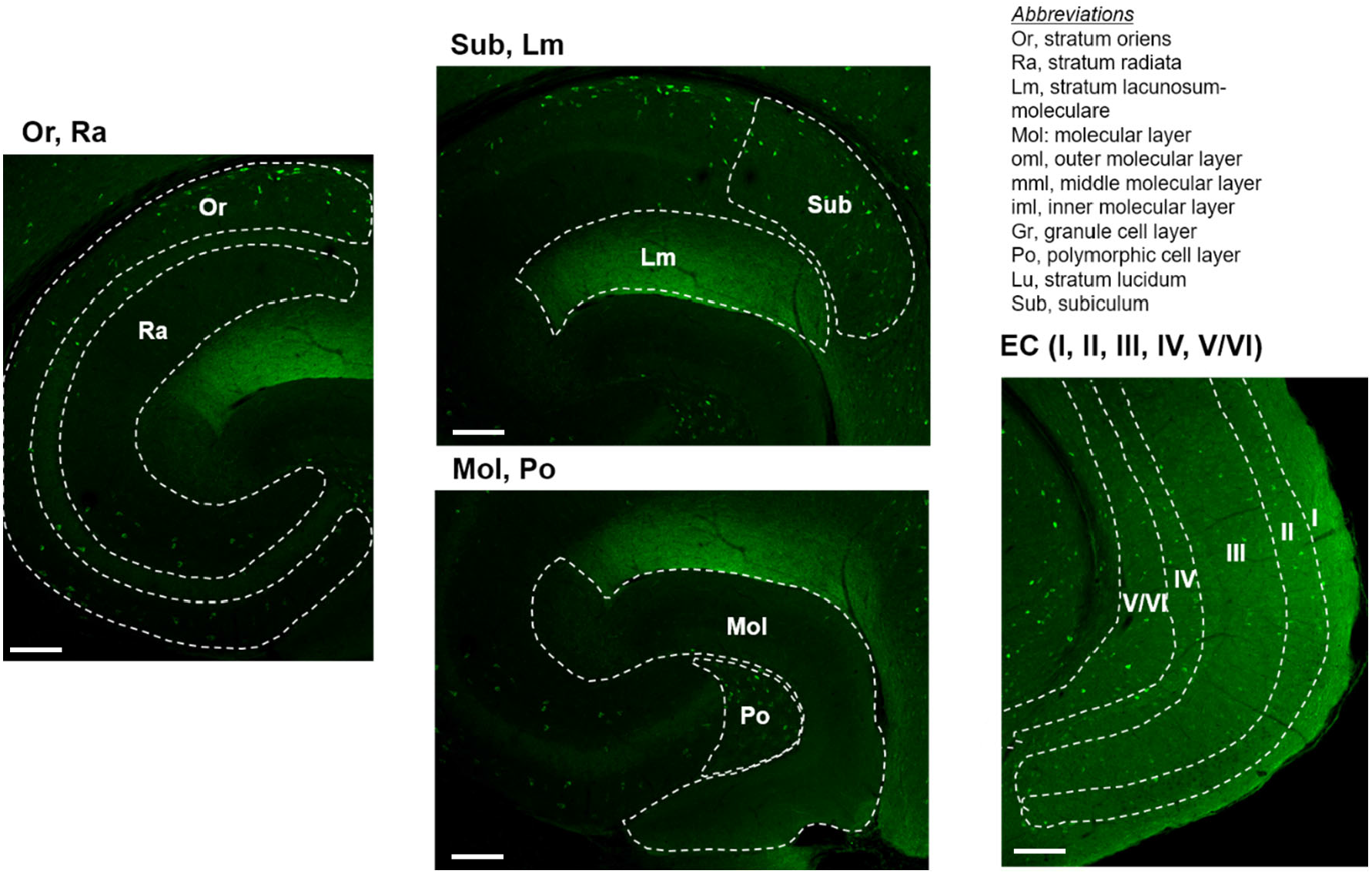
Immunohistochemistry of somatostatin in the hippocampus. The upper right panel defines abbreviations used in the images. The specificity of immunostaining was confirmed using mice deficient in somatostatin precursor as a negative control. The scale bar indicates 100 μm.

**Fig. 3S.**
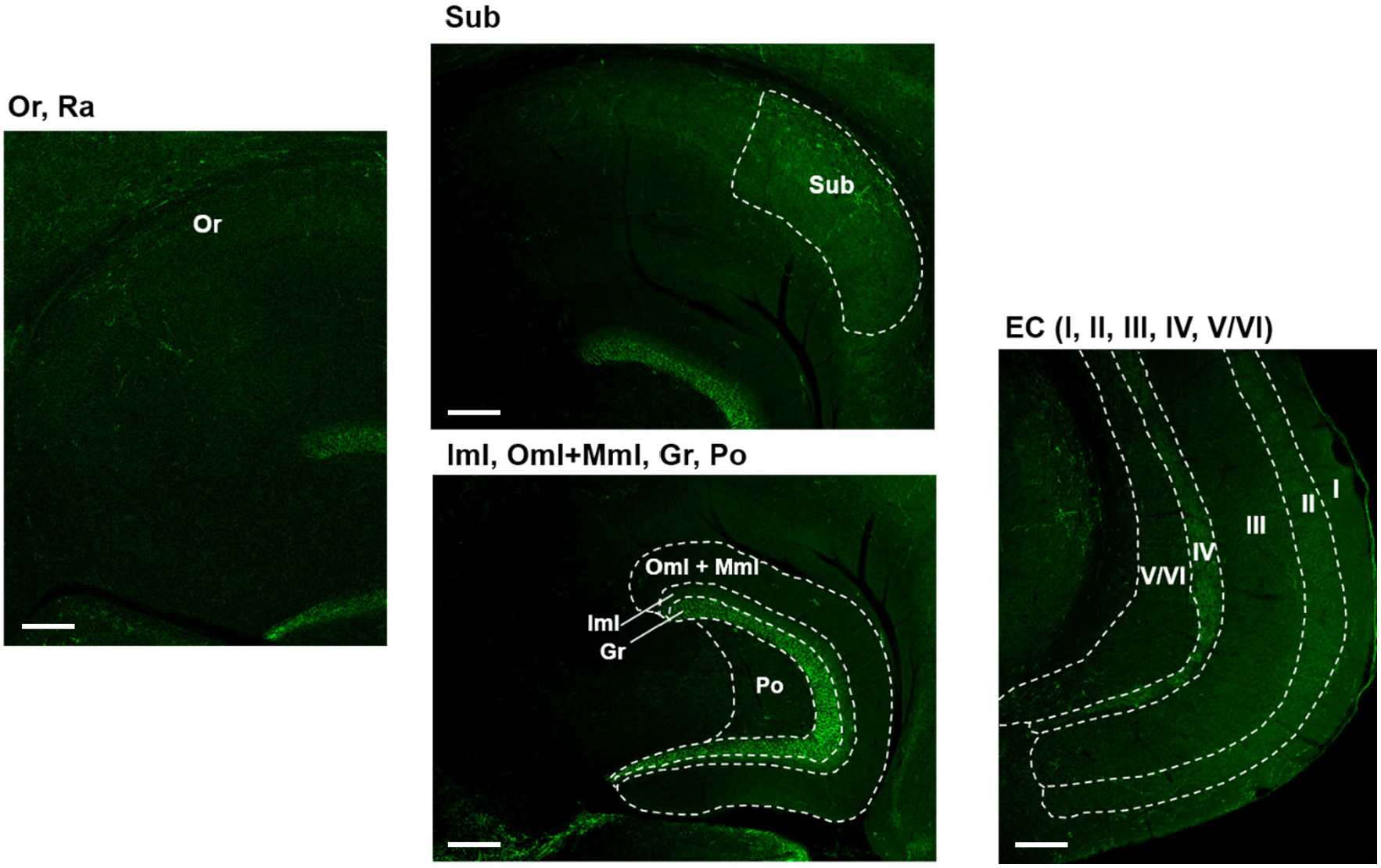
Immunohistochemistry of substance P in the hippocampus. Abbreviations used in the images are defined in Fig. 2S. The scale bar indicates 100 μ m.

**Fig. 4S.**
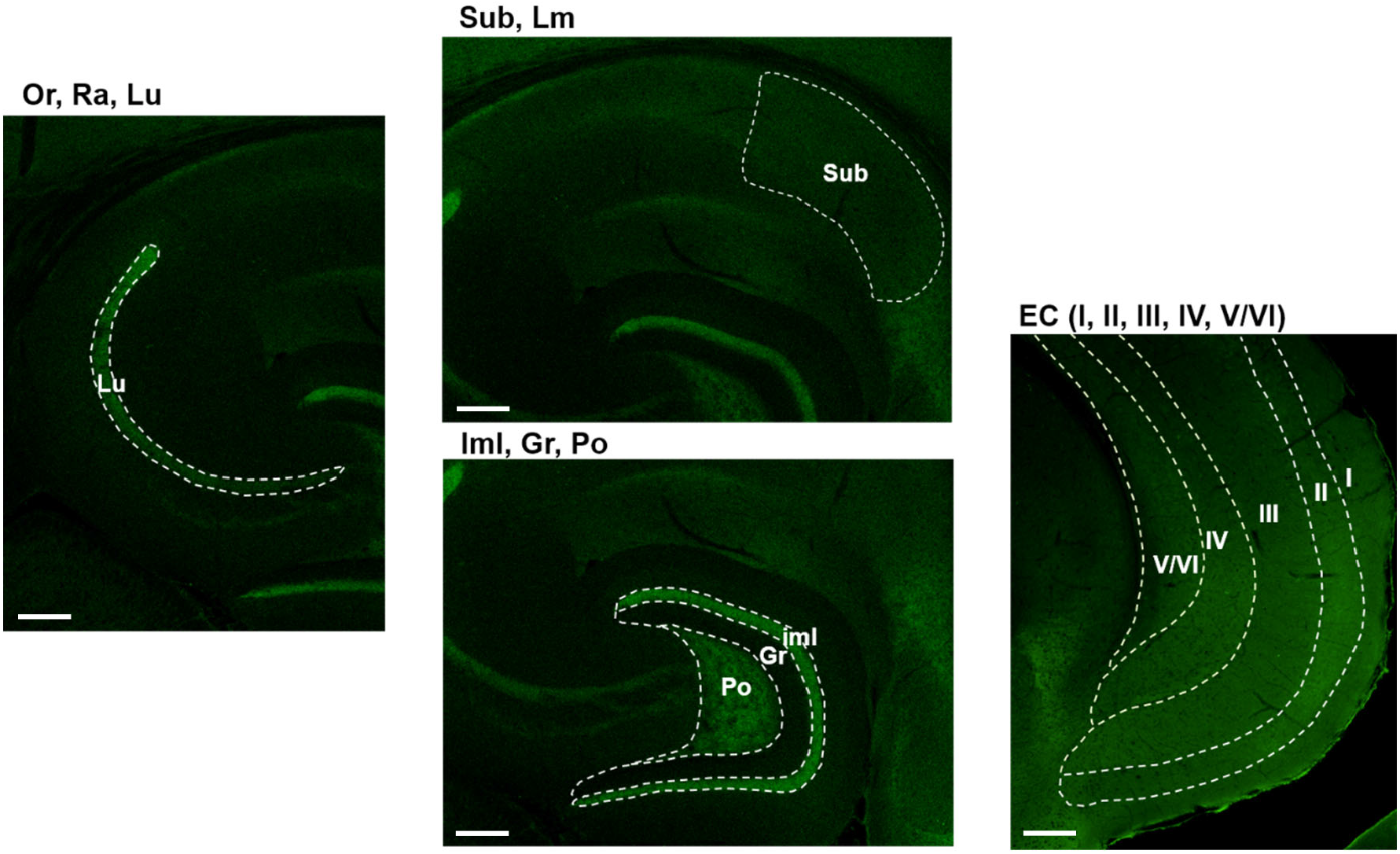
Immunohistochemistry of cholecystokinin in the hippocampus. Abbreviations used in the images are defined in Fig. 2S. The scale bar indicates 100 μm.

**Fig. 5S.**
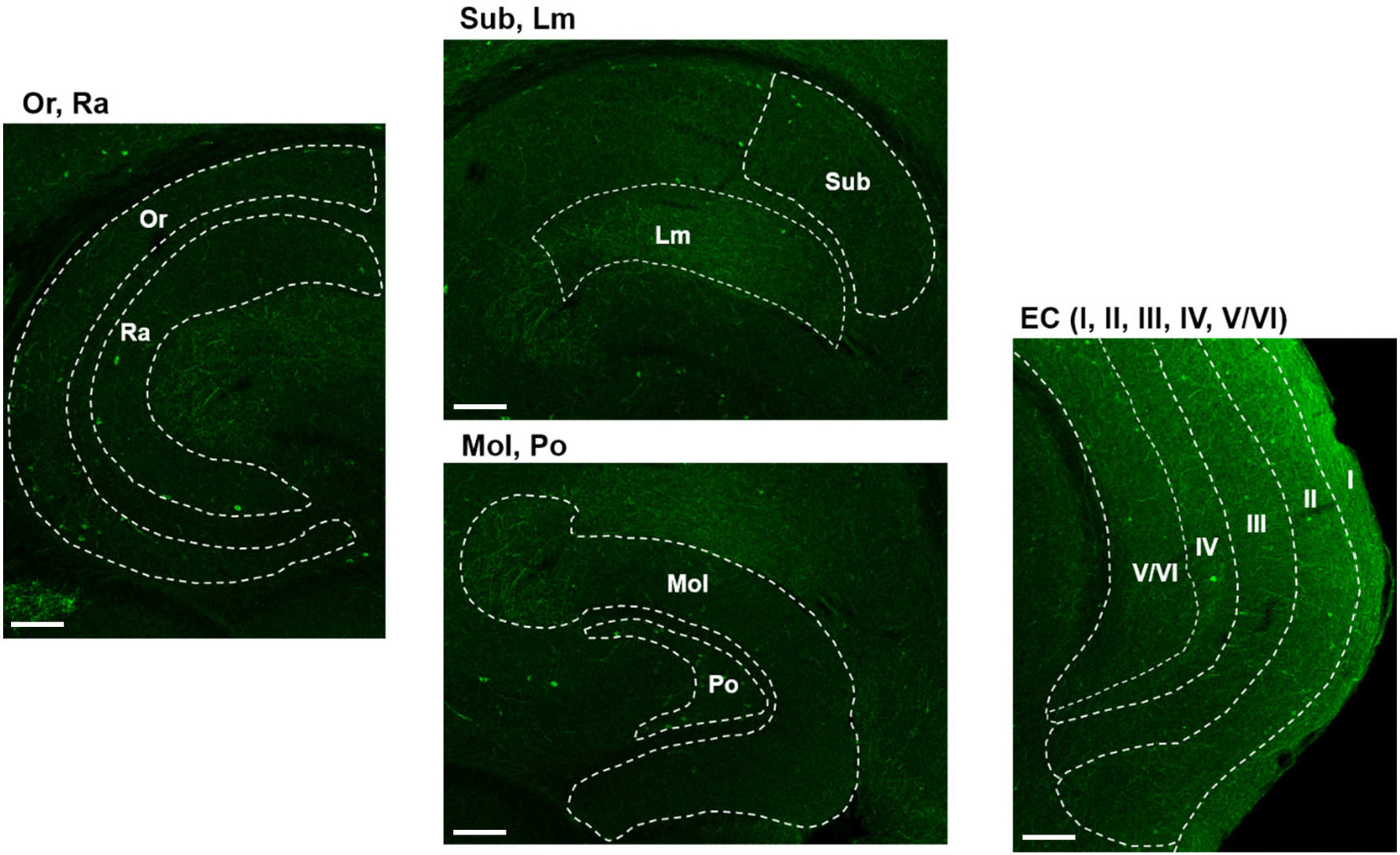
Immunohistochemistry of neuropeptide Y in the hippocampus. Abbreviations used in the images are defined in Fig. 2S. The scale bar indicates 100 μm.

**Fig. 6S.**
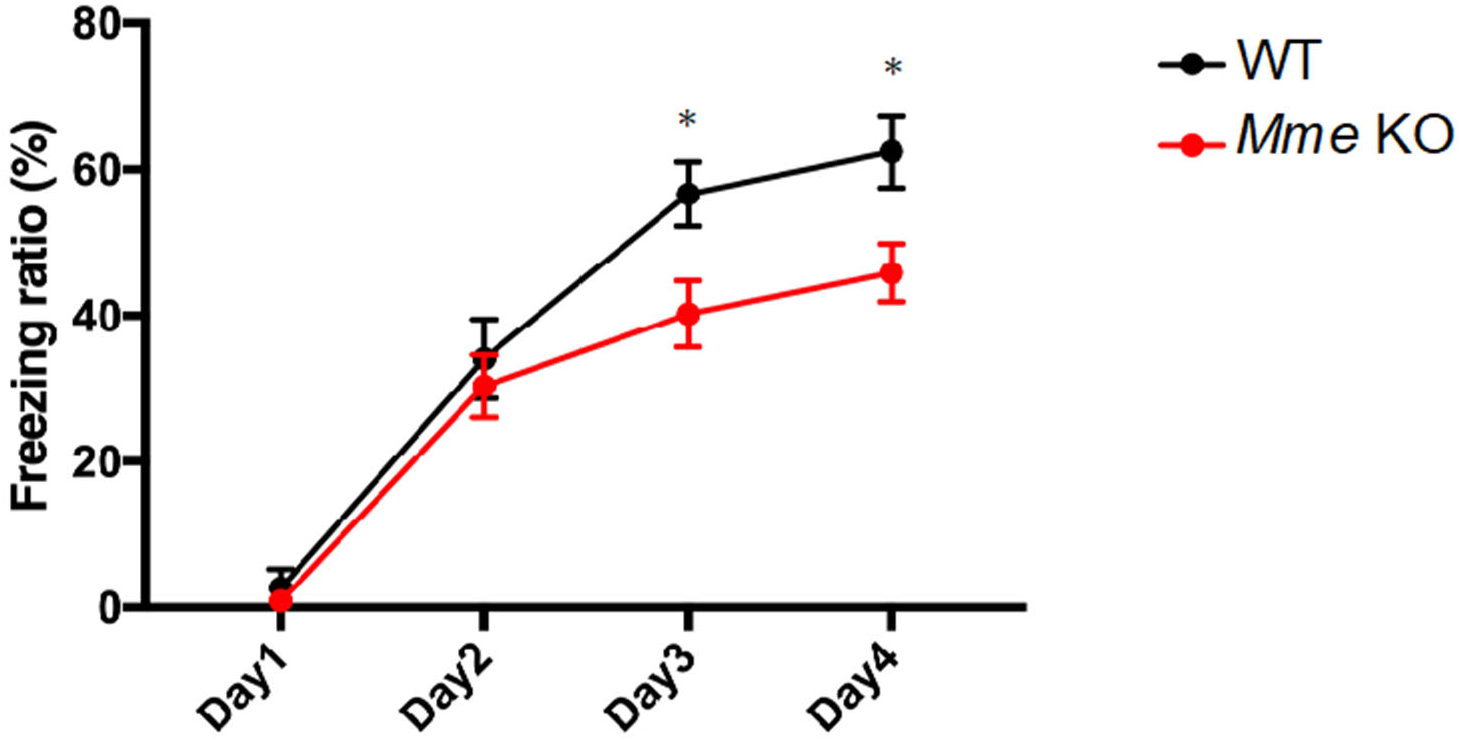
Memory impairment in single NEP (*Mme*) KO mice as analyzed by fear conditioning. Freezing ratio of 6-month-old WT and NEP-KO mice. The data represent the mean ±SEM. Results were analyzed by two-way ANOVA followed by Bonferroni’s multiple comparison test (n = 8 for each group).

## Supplementary Methods

### Gel filtration

TS-soluble fractions of mouse cerebral cortex (10.6mg) were applied to a gel filtration column (TSK gel G2000SWXL, 7.8×300mm) (TOSOH, Tokyo, Japan) equilibrated with buffer A (50% acetonitrile-0.1% trifluoroacetic acid). The adsorbed material was eluted with buffer A at a flow rate of 1ml/min at 40°C. Fractions (500μl) were pooled, lyophilized, and subjected to ELISA for Aβ_40_ and Aβ_42_ quantification. The elution volumes of Aβ monomer and oligomers were determined using a monomer and oligomers generated from synthetic Aβ. Aβ_42_ (PEPTIDE INSTITUTE. INC., Osaka, Japan) (2mg/ml DMSO, 10μl) was added to 20mM Tris-HCl (pH 7.5, 90μl) and then incubated for 1 day at 37°C.

### Immunohistochemical staining and image analysis of neuropeptides in WT and NEP-KO mouse hippocampi

Male neprilysin-knockout mice/B6J (12-14 weeks of age; n=7) and wild type littermates (n=8) were used. The mice were anesthetized deeply and perfused transcardially with 0.1 M PBS (pH 7.4), followed by ice-cold Zamboni’s fixative (+1.5% glutaradehyde). Brains were removed and immersed in the same fixative at 4°C overnight. Horizontal sections (40 μm thick) were prepared using a vibrating blade microtome (VT1000 S vibratome, Leica Biosystems), and were immunostained floating in the primary antibody solution for overnight. Primary antibodies against neuropeptides and secondary antibodies used were as follows ^1-4^: anti-somatostatin-14 (rabbit polyclonal, 1:4000 dilution, Peninsula Laboratories Cat# T-4103.0050, RRID:AB_518614), anti-substance P (rat monoclonal, 1:50 dilution, Millipore Cat# MAB356, RRID:AB_94639), anti-neuropeptide Y (rabbit polyclonal, 1:6000 dilution, Sigma-Aldrich Cat# N9528, RRID:AB_260814), anti-cholecystokinin 8 (rabbit polyclonal, #20078 1:6000 dilution, Diasorin, Stillwater, MN), AlexaFluor 488-conjugated anti-mouse and anti-rat antibody (1:500 dilution, Thermo Fisher Scientific Cat# A-11034, RRID:AB_2576217, Thermo Fisher Scientific Cat# A-11006, RRID:AB_2534074). Confocal images were acquired by a triple scan protocol with an IX70 inverted microscope incorporating an FV300 confocal laser scanning system. The densities of immunoreactivity to each antibody in grid areas of the hippocampal formation and entorhinal cortex were measured using MetaMorph, ver. 7.7 (Molecular Devices). To reduce the variance of tissue sections, we used the average of data from 9-10 tissue sections per mouse as an individual value. In addition, each set of experiments was repeated at least twice to confirm the results.

*Note*. Changes in levels of Met-Enkephalin, Leu-Enkephalin, CRF (Corticotropin-releasing factor), Neurotensin B and other neuropeptides were not examined because of their low expression levels in the hippocampal formation and entorhinal cortex.

### Statistical analysis

All data for immunohistochemical images were expressed as means ± s.e.m. For comparisons of the means between two groups, statistical analysis was performed by applying Student’s *t* test after confirming equality of variances of the groups. If the variances were unequal, a Mann-Whitney *U*-test was performed (SigmaPlot software, ver.14, Systat Software Inc).

### Examination of memory impairment by fear conditioning

We performed the memory impairment examination as previously described^5^. Before the start of test, mice were put in a white noise box for at least 1 hour. Subsequently, the mice were placed into a sound-attenuating chamber and allowed to explore the chamber for 5 minutes. The percent freezing time was measured until mice received an electric shock (7.5mA) to the foot after 4 minutes. As a long-term retention test, the same conditioning experiments were repeated daily for 4 days. The training box was cleaned with water and wiped dry with paper toweling before the next mouse was placed in the chamber. Mice were returned to their cages and provided with free access to food and water.

## Notes

### Competing Interest Statement

The authors have declared no competing interest.

